# mosdepth: quick coverage calculation for genomes and exomes

**DOI:** 10.1101/185843

**Authors:** Brent S. Pedersen, Aaron R. Quinlan

**Affiliations:** University of Utah, Department of Human Genetics, Department of Biomedical Informatics, and USTAR Center for Genetic Discovery.

## Abstract

*Mosdepth* is a new command-line tool for rapidly calculating genome-wide sequencing coverage. It measures depth from BAM (Li et al. 2009) or CRAM files at either each nucleotide position in a genome or for sets of genomic regions. Genomic regions may be specified as either a BED file to evaluate coverage across capture regions, or as a fixed-size window as required for copy-number calling. Mosdepth uses a simple algorithm that is computationally efficient and enables it to quickly produce optional coverage summaries. We demonstrate that *mosdepth* is faster than existing tools and provides flexibility in the types of coverage profiles produced.

**Availability:** *mosdepth* is available from https://github.com/brentp/mosdepth under the MIT license‥

**Supplementary information:** Detailed documentation is available at https://github.com/brentp/mosdepth

## 1 Introduction

Measuring the depth of sequencing coverage is critical for genomic analyses such as calling copy-number variation (CNV), e.g., by *cn.mops* (Klambauer *et al.*, 2012), quality control (Pedersen *et al.*, 2017), and determining which genomic regions have too low, or too high (Li, 2014) coverage for reliable variant calling. Given the scope of applications for coverage profiles, there are several existing tools that calculate genome-wide coverage. Samtools depth (Li *et al.*, 2009) outputs per-base coverage; BEDTools genomecov (Quinlan and Hall, 2010; Quinlan, 2014) can output per-region or per-base coverage; Sambamba (Tarasov *et al.*, 2015) also provides per-base and per-window depth calculations. The need for efficient coverage calculation increases with the number and depth of whole genome sequences, and existing methods require roughly an hour or more of computation for a typical human genome with 30x coverage. Here, we compare *mosdepth* to these existing methods.

## 2 Methods

*Mosdepth* uses HTSLib (http://www.htslib.org/) via the nim programming language (https://nim-lang.org); it expects the input BAM or CRAM file to be sorted by position. In contrast to samtools, which uses a “pileup” engine that tracks each nucleotide in every read, *mosdepth* only tracks chunks of read alignments. Only the start and end position of each chunk of an alignment (each alignment may have multiple chunks if it is split by a deletion or other event) are tracked in an array (of 32 bit integers) whose sizeis the length ofthe chromosome. For each chunkof an alignmentto the reference genome, *mosdepth* increments the start and decrements the end forthe the valueatthe indexinthe array correspondingtothat chromosomal position (Figure 1). It avoids double-counting coverage when the ends of a paired-end sequencing fragment have overlapping alignments (Figure 1, black alignment). Once the coverage array has tracked all alignment starts and ends in a BAM or CRAM file, the depth at a particular position is calculated as the cumulative sum of all array positions preceding it (a similar algorithm is used in BEDTools which track starts and ends separately).

**Fig. 1.**
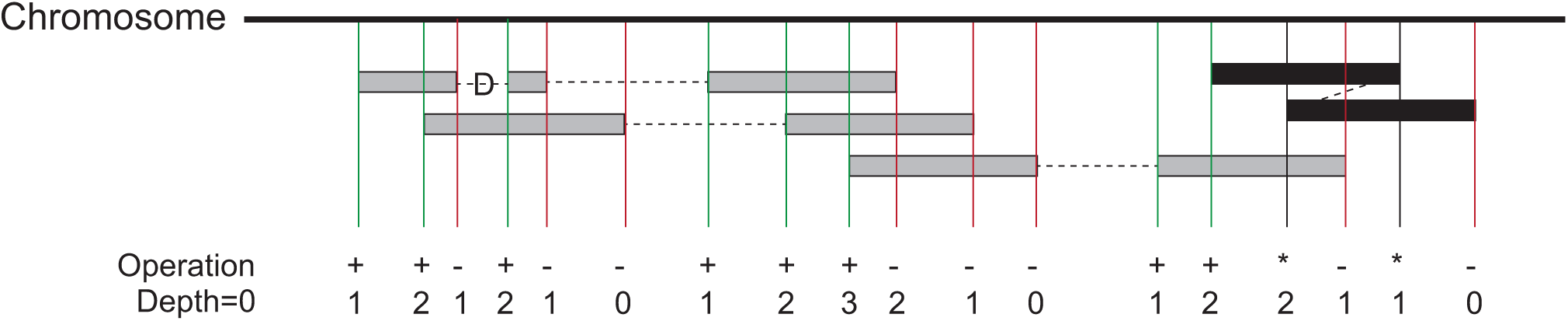
*Mosdepth* coverage calculation algorithm. An array the size of the current chromosome is allocated. As each alignment is read from a position-sorted BAM or CRAM file, the value at each start is incremented and the value at each stop is decremented. As illustrated by the alignment with a deletion (D) CIGAR operation, each alignment may have multiple starts and ends. If the leftmost read (the one seen first) of a paired-end alignment has an end that overlaps the position of its mate (which is given as a field in the BAM record) then it is stored in a hash-table until its mate is seen. At that time, the overlap between the mates is calculated, the regions of overlap are decremented and the item is removed from the hash. This prevents double counting coverage from two ends of the same paired-end DNA fragement (black alignment, “*” operation means no coverage increment or decrement is made). Once all reads for a chromosome are consumed, the per-base coverage is simply the cumulative sum of the preceding positions.

The coverage along a chromosome is calculated in place by replacing the composite start and end counts with the cumulative sum up to each element in the array. Once complete, the coverage of a region is simply the mean of the elements in the array spanning from start to end. This makes it possible to calculate coverage extremely quickly, even for millions of small regions. This setup is also amenable to rapid calculation of a genome’s coverage distribution: that is, the number of bases covered by a given number of reads across the genome or in the given regions. The distribution calculation requires an extra iteration through the array that counts the occurrence of each coverage value. The *mosdepth* method does require more memory–for the 249 megabase chromosome 1 in the human genome, it will require about 500MB of memory plus additional because of delay in garbage-collection, however, that number is not dependent on the depth of coverage or number of alignments. Despite its flexibility, *mosdepth* is easy to use and understand (see Supplement for example uses).

## 3 Results

We compared the time and memory requirements of *mosdepth* (v0.1.6) to samtools (v1.5), BEDTools (v2.26.0) and sambamba (v0.6.6) on a BAM with about 30X coverage from the Simon’s Genome Diversity Panel (Mallick *et al.*, 2016) (Supplemental Info). With a single CPU, mosdepth is faster than existing tools, and can be even faster with multiple threads (Table 1). Results for CRAM and for other options such as window-based depth calculations are shown in Supplemental Table 1.

**Table 1.**
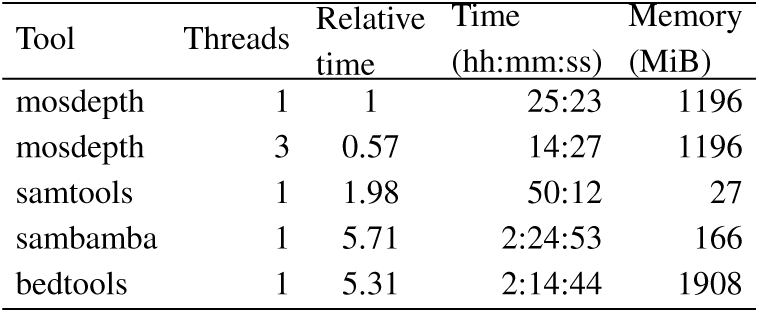
Comparison of depth tools for time and memory use on a 30X BAM. Mosdepth and BEDTools use much more memory, but mosdepth isnearly twice as fast as the next fastest tool, samtools.

| Tool | Threads | Relative time | Time (hh:mm:ss) | Memory (MiB) |
| --- | --- | --- | --- | --- |
| mosdepth | 1 | 1 | 25:23 | 1196 |
| mosdepth | 3 | 0.57 | 14:27 | 1196 |
| samtools | 1 | 1.98 | 50:12 | 27 |
| sambamba | 1 | 5.71 | 2:24:53 | 166 |
| bedtools | 1 | 5.31 | 2:14:44 | 1908 |

To evaluate consistency between the tools, we compared the output to samtools depth. *Mosdepth* cannot include or exclude individual bases because of low base-quality (BQ) as can samtools depth. In contrast, samtools depth cannot avoid double-counting overlapping regions unless the BQ cutoff is set to a value > 0. Therefore, we compared *mosdepth* without mate overlap correction to samtools depth with a BQ cutoff of 0 for chromosome 22 of the dataset used for Table 1. With this comparison set up to evaluate differences, we found no discrepancies in reported depth among the tools for the entire chromosome.

## 4 Discussion

*Mosdepth* is a quick, convenient tool for genome-wide depth calculation. The optional coverage distribution is useful for quality control and the depth output is applicable without further processingasinputtomany CNV detection tools. While the method it employs requires greater memory use, it makes the implementation simple and fast, enables a straightforward coverage distribution calculation, and expedites the depth calculations for even millions of regions. *Mosdepth* is useful for exome, whole-genome, and targeted sequencing projects.

